# Recombinant spore-displayed enzymes for cellulose hydrolysis: dual-enzyme spores versus mixed single-enzyme spores

**DOI:** 10.64898/2026.10.21.760755

**Authors:** Rohit P. James, Santosh B. Noronha

## Abstract

*Bacillus subtilis* spore surface display has been widely used to immobilise single recombinant enzymes, yet many industrially relevant reactions are multi-enzyme cascades. This study examined whether two distinct, catalytically active enzymes can be co-displayed on a single spore, using the two-step hydrolysis of carboxymethyl cellulose (CMC) to glucose as a model cascade. An endoglucanase (EglS) and a *β*-glucosidase (Unbgl1A) were expressed in *E. coli*, characterised, and surface-displayed on *Bacillus subtilis* WB800N by fusion to the coat anchors CotB, CotC and CotG. The best constructs were combined either by mixing two single-enzyme spore strains or by co-displaying both enzymes on one strain, and compared with the free-enzyme mixture on CMC. CotG was the most effective anchor for EglS and the only functional anchor for Unbgl1A, with improved thermal stability relative to the free enzymes and activity retained over repeated reuse: CotG-EglS retained ~ 46% activity after 12 h (versus ~ 27% for the free enzyme), and CotG-Unbgl1A retained ~ 30% activity after 8 h, whereas free Unbgl1A lost all activity within 6 h. The mixed single-enzyme spores gave the highest glucose yield (2.15 mM), exceeding both the single dual-enzyme strain (1.57 mM) and the free-enzyme mix (1.35 mM). Two functional enzymes were co-displayed on one spore, but mixing separate single-enzyme spores gave higher yield under the tested conditions and allowed the enzyme ratio to be tuned.

## 1 Introduction

Spore surface display is a technique in which a protein of interest is displayed on the surface of spores by genetic fusion with native surface coat proteins. *Bacillus subtilis*, a model sporulating Gram-positive bacterium, has emerged as an established chassis for spore surface display (Iwanicki et al., 2014). The native proteins of the outer coat and crust layers are fused in frame with a passenger enzyme via a linker intended to facilitate independent folding, and the fusion is expressed under a sporulation-specific promoter so that it integrates during spore coat assembly. The coat proteins CotB, CotC and CotG are among the most frequently used anchors and were chosen for this study. Spore surface display has been reported to improve the thermal stability (Wang et al., 2017) and recyclability (Lehmann and Herrmann, 2025) of enzymes, and is thus a natural method of enzyme immobilisation.

The robustness of *Bacillus subtilis* spore surface display has led to diverse applications ranging from vaccine development (Nguyen et al., 2025) to biosensors. However, these applications have relied on the surface display of a single recombinant passenger per spore, typically an antigen or an enzyme, whereas many industrially relevant reactions involve multiple enzymes. To our knowledge, the co-display of two distinct, catalytically active enzymes on a single spore surface has not previously been reported. To assess the feasibility of spore surface display for multi-enzyme cascades, a model two-step enzymatic cascade was used: the hydrolysis of carboxymethyl cellulose (CMC) by spore-displayed cellulases. Industrial cellulose hydrolysis relies on multi-component cellulase cocktails, but a minimal route to glucose requires only two activities, an endoglucanase and a *β*-glucosidase (Nath et al., 2019). Two enzymes were therefore used: an endoglucanase (EglS), which cleaves internal *β*-1,4-glycosidic linkages in the cellulose polymer to release chain ends and small oligosaccharides such as cellobiose, and a *β*-glucosidase (Unbgl1A), which breaks down these oligosaccharides into glucose.

This study aimed to determine whether dual-enzyme surface display on *Bacillus subtilis* spores is feasible. A two-step, cascading enzymatic reaction was necessary to evaluate the function of both displayed enzymes, both independently and synergistically. The two enzymes were first expressed in *E. coli* and characterised, then cloned into the vectors pBS1C and pBS4S as fusions to CotB, CotC and CotG and displayed on *Bacillus subtilis* WB800N. The best single-enzyme constructs were then combined in two ways, by mixing separate single-enzyme spore strains and by co-displaying both enzymes on one strain, and their performance on CMC was compared with that of the free-enzyme mixture.

## 2 Materials and methods

### 2.1 Strains, plasmids and cloning

Cloning was performed in *E. coli* DH5*α*, and proteins were expressed in *E. coli* BL21(DE3) using the pET43 vector. Expression of EglS and Unbgl1A was induced with 0.15 mM IPTG, followed by incubation at 25°C and 120 rpm for 8 h. A His-tag was integrated at the C-terminus through primers during template amplification and used for purification from cell lysate with Ni-NTA agarose beads. EglS (~1413 bp) and Unbgl1A (~ 1410 bp) were each cloned into pET43. For surface display, the coat genes *cotB, cotC* and *cotG* were amplified together with their native promoters from *Bacillus subtilis* 168 genomic DNA and cloned into the integrative shuttle vectors pBS1C and pBS4S (Radeck et al., 2013); a flexible G-G-G-G-S linker was incorporated between the anchor and the passenger. The cot partner was cloned between EcoRI and XbaI, and the passenger between XbaI and PstI (pBS1C) or XbaI and NotI (pBS4S). The constructs were transformed into *Bacillus subtilis* WB800N by the two-step natural-competence protocol (Harwood and Cutting, 1990), generating six strains: CotB-EglS, CotC-EglS, CotG-EglS, CotB-Unbgl1A, CotC-Unbgl1A and CotG-Unbgl1A. The dual-enzyme strain was generated by integrating CotG-EglS (pBS1C) as a single chromosomal copy at the *amyE* locus and CotG-Unbgl1A (pBS4S) as a single chromosomal copy at the *thrC* locus in one *Bacillus subtilis* WB800N isolate, so that both fusion cassettes were stably maintained in every cell of the resulting clonal strain. Selection used ampicillin for *E. coli*; in *Bacillus subtilis*, the EglS constructs (pBS1C) were selected on chloramphenicol and the Unbgl1A constructs (pBS4S) on spectinomycin, with the dual-enzyme strain selected on both antibiotics.

### 2.2 Sporulation

Sporulation was induced by the exhaustion method (Harwood and Cutting, 1990). Cells were grown in sporulation medium for 36–48 h with periodic microscopy until more than 90% had sporulated. Spores were harvested by centrifugation, washed with 0.5 M NaCl followed by 1 M KCl, resuspended in 50 mM Tris-Cl (pH 7.2), treated with 1 mg/mL lysozyme for 1 h at 4°C to lyse surviving vegetative cells, washed once with 1 M NaCl and thrice with distilled water, and stored at 4°C.

### 2.3 Enzyme assays

Endoglucanase (EglS) activity was characterised using 0.5% carboxymethyl cellulose as substrate, with reducing sugars quantified by the DNSA method (Rahman et al., 2018). One unit (U) of endoglucanase activity was defined as the amount of enzyme required to release 1 *µ*mol of reducing sugars per minute from CMC under optimal assay conditions. *β*-glucosidase (Unbgl1A) activity was characterised separately on 15 mM cellobiose, with glucose quantified by the GOD-POD method (Borges et al., 2014). One unit (U) of *β*-glucosidase activity was defined as the amount of enzyme releasing 1 *µ*mol of glucose per minute under the specified assay conditions. Assays were performed in 50 mM sodium acetate buffer.

### 2.4 Surface-display confirmation and enzyme loading

Surface accessibility was probed by a protease accessibility test using pepsin and trypsin, each at 1 mg/mL, at 37°C for 1 h; a loss of activity relative to the untreated control indicates that the displayed enzyme is accessible at the spore surface. For loading, recombinant spores were assayed under optimal conditions, and the measured activity of each construct was converted to the corresponding amount of functional enzyme per 10^10^ spores using the specific activity of the respective purified free enzyme. This activity-based estimate reflects the functionally active displayed enzyme, assuming comparable specific activity for the displayed and free forms, rather than total displayed protein.

### 2.5 Optima, thermal stability and recycling

Spore-displayed and free enzymes were compared across pH and temperature to determine their optima. The pH profiles were determined at a fixed temperature of 45°C, and the temperature profiles were determined at a fixed pH of 7. Thermal stability was assessed from residual activity during incubation at 45°C for EglS (12 h) and 50°C for Unbgl1A (8 h). For recycling, spores were recovered by centrifugation after each reaction, washed, and reused. Activity in the first use, shown as Control for EglS and 0 for Unbgl1A, was taken as 100%; rounds 1–4 indicate subsequent reuses.

### 2.6 Two-enzyme hydrolysis of CMC

The operating pH, temperature and endoglucanase:*β*-glucosidase activity ratio were optimised for sequential CMC hydrolysis (see Results). The cascade was then run with 1% CMC in 3 mL at pH 5 and 50°C, and glucose release was quantified by the GOD-POD method over 8 h. Before mixing, EglS and Unbgl1A activities were standardised separately using 1% CMC and 15 mM cellobiose, respectively. The mixed single-enzyme spores and the free-enzyme mixture each contained 0.5 U EglS and 0.25 U Unbgl1A.

The dual-enzyme strain was dosed to provide the same 0.5 U EglS, which supplied 0.3 U Unbgl1A because the two activities could not be adjusted independently. An initial experiment compared mixed single-enzyme spores with the free-enzyme mixture and individual-enzyme controls. After construction of the dual-enzyme strain, a separate experiment compared all three preparations. Starting EglS activity was matched in both experiments.

### 2.7 Statistical analysis

All experiments were performed in triplicate (n = 3 independent biological replicates). Data are expressed as mean ± standard deviation. Results were plotted using OriginPro 2022b.

## 3 Results

### 3.1 Constructs, expression and surface display

EglS and Unbgl1A were cloned into pET43 (Figs. 1 and 2) and expressed in *E. coli* BL21(DE3) under IPTG induction. SDS-PAGE confirmed expression of EglS at its theoretical size of 50 kDa and of Unbgl1A at 55 kDa, against an empty-vector NusA-tag control (Figs. 3 and 4). The two enzymes were then fused to CotB, CotC and CotG and displayed on *Bacillus subtilis* WB800N (Table 1); restriction analyses of the EglS and CotG-Unbgl1A display constructs are shown in Figs. 5 and 6, respectively. Among the six fusion strains, CotB-Unbgl1A and CotC-Unbgl1A were non-functional, leaving CotG-Unbgl1A as the only functional spore-displayed *β*-glucosidase strain, whereas all three EglS constructs were functional.

**Table 1.** Strains used in this study.

| Strain | Description |
| --- | --- |
| pET43-EglS (*E. coli*) | Referred to as free enzyme EglS |
| pET43-Unbgl1A (*E. coli*) | Referred to as free enzyme Unbgl1A |
| CotB-EglS (*B. subtilis* spore) | Fusion of coat protein CotB and EglS displayed on the spore surface |
| CotC-EglS (*B. subtilis* spore) | Fusion of coat protein CotC and EglS displayed on the spore surface |
| CotG-EglS (*B. subtilis* spore) | Fusion of coat protein CotG and EglS displayed on the spore surface |
| CotG-Unbgl1A (*B. subtilis* spore) | Fusion of coat protein CotG and Unbgl1A displayed on the spore surface |
| Dual strain (CotG-EglS + CotG-Unbgl1A) | Single strain co-displaying both enzymes; cassettes integrated at *amyE* (pBS1C) and *thrC* (pBS4S) |

**Figure 1.**
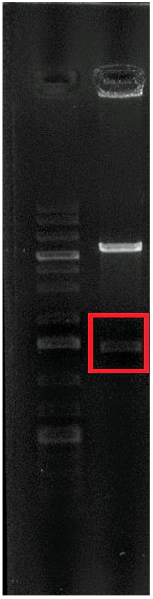
Restriction analysis of pET43-EglS.

**Figure 2.**
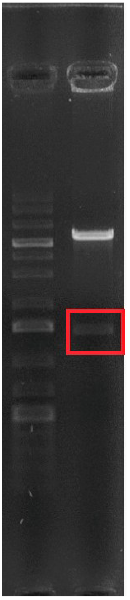
Restriction analysis of pET43-Unbgl1A.

**Figure 3.**
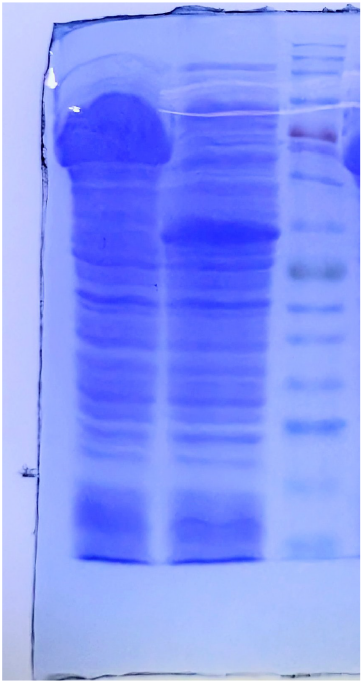
SDS-PAGE of pET43-EglS. Lanes: 1, empty pET43-NusA control; 2, EglS under IPTG induction; 3, protein ladder.

**Figure 4.**
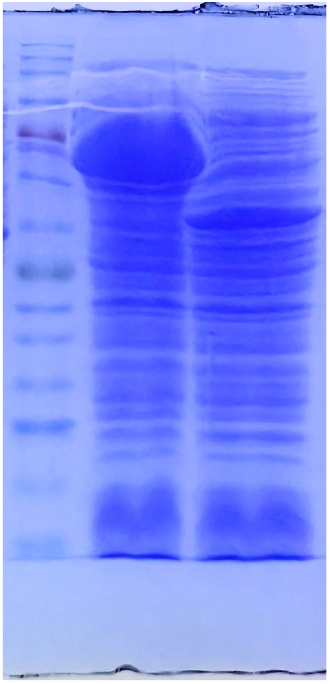
SDS-PAGE of pET43-Unbgl1A. Lanes: 1, protein ladder; 2, empty pET43-NusA control; 3, Unbgl1A under IPTG induction.

**Figure 5.**
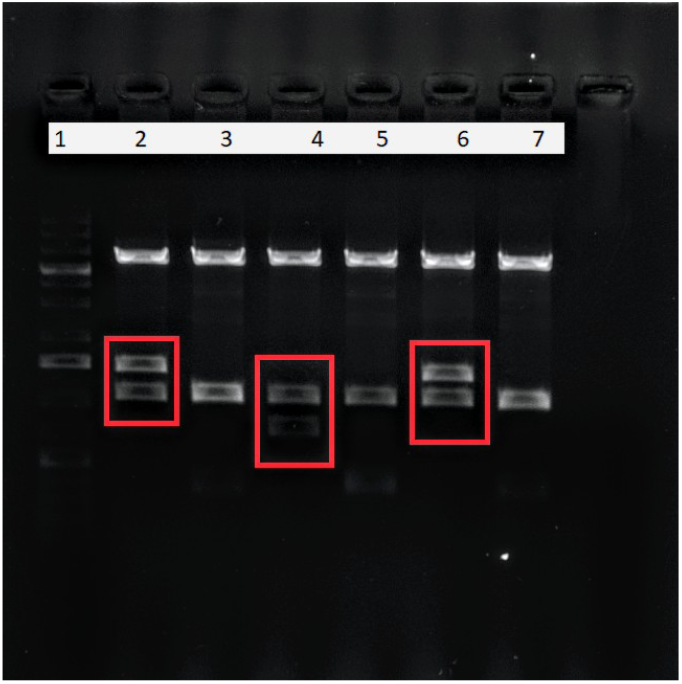
Restriction analysis of the pBS1C cot-EglS constructs in the order CotB-EglS, CotC-EglS and CotG-EglS.

**Figure 6.**
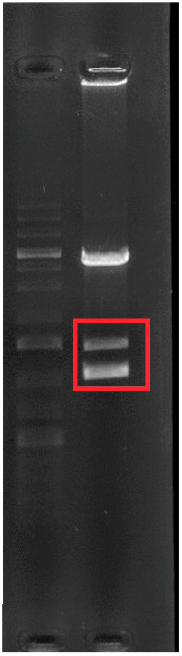
Restriction analysis of the pBS4S-CotG-Unbgl1A construct.

All surface-displayed EglS constructs showed a loss of activity on treatment with pepsin and trypsin, indicating that the displayed enzyme is accessible at the spore surface; the loss was greater with trypsin, and CotG-EglS retained the least activity, consistent with it being the most surface-exposed construct (Fig. 7). CotG-Unbgl1A likewise lost activity on protease treatment, indicating its surface accessibility (Fig. 8).

**Figure 7.**
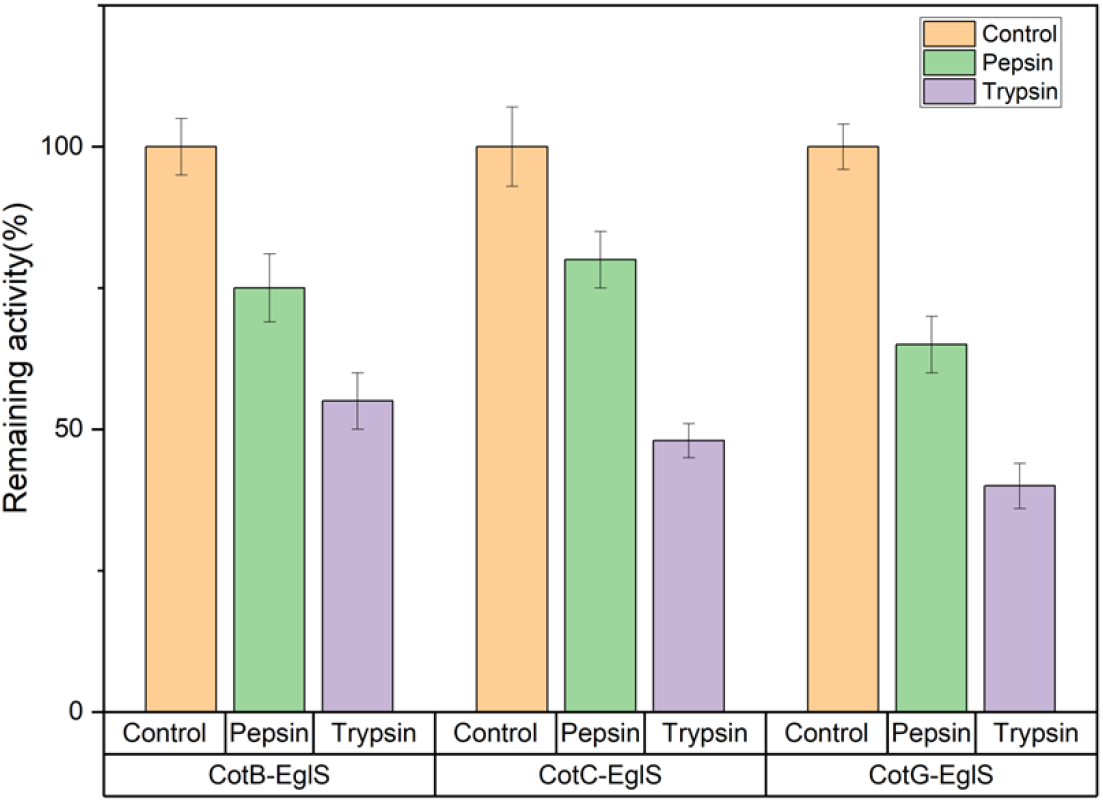
Effect of pepsin and trypsin on spore-displayed EglS after 1 h at 37°C. Activity is expressed relative to the untreated control (100%). Quantitative plots in this paper show mean ± SD from three independent biological replicates (n = 3).

**Figure 8.**
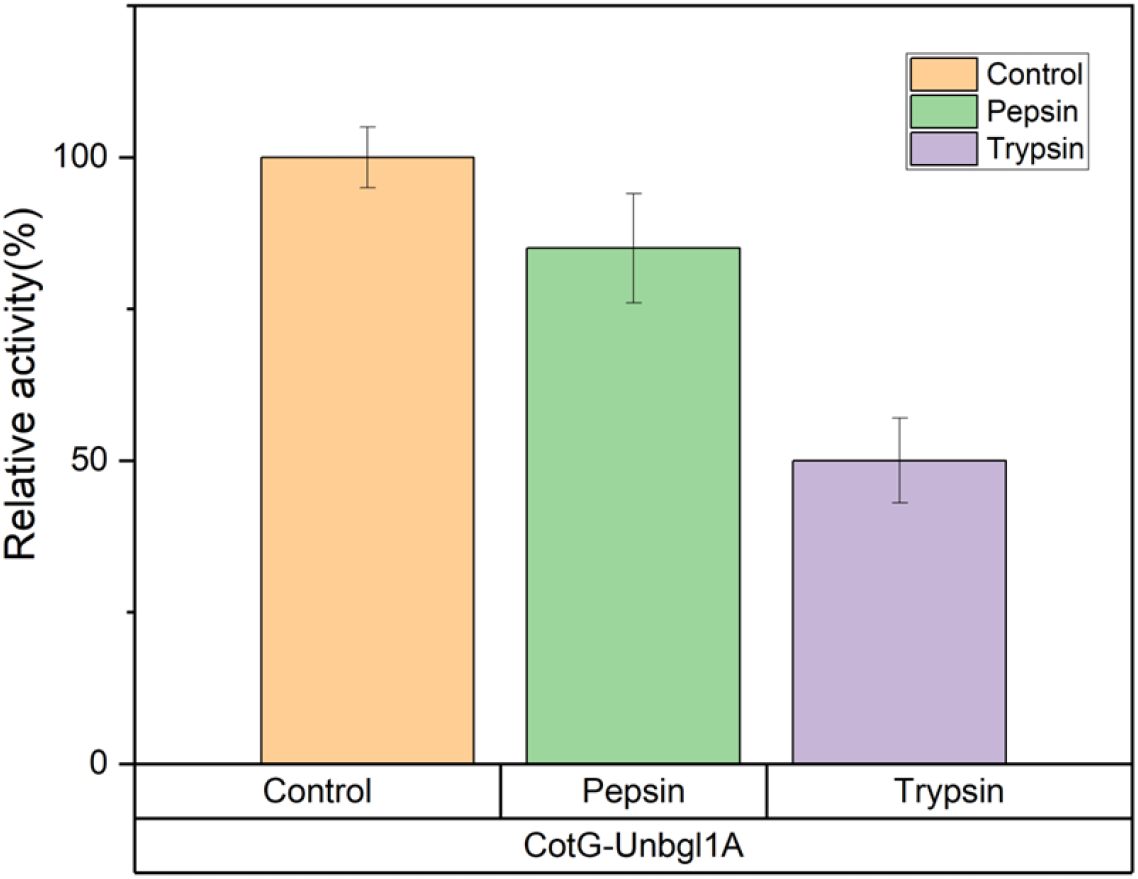
Effect of pepsin and trypsin on CotG-Unbgl1A relative to the untreated control.

The functional loading of each construct was estimated from its activity (Fig. 9). Among the EglS constructs, CotG-EglS showed the highest functional loading at 2.5 *µ*g per 10^10^ spores, followed by CotB-EglS (2.0 *µ*g) and CotC-EglS (1.6 *µ*g). The single-enzyme CotG-Unbgl1A construct loaded 4.0 *µ*g per 10^10^ spores. In the dual-enzyme strain, loading was lower for both enzymes (1.4 *µ*g for EglS and 2.0 *µ*g for Unbgl1A per 10^10^ spores), consistent with competition between the two CotG fusions and the native CotG for incorporation into the spore coat.

**Figure 9.**
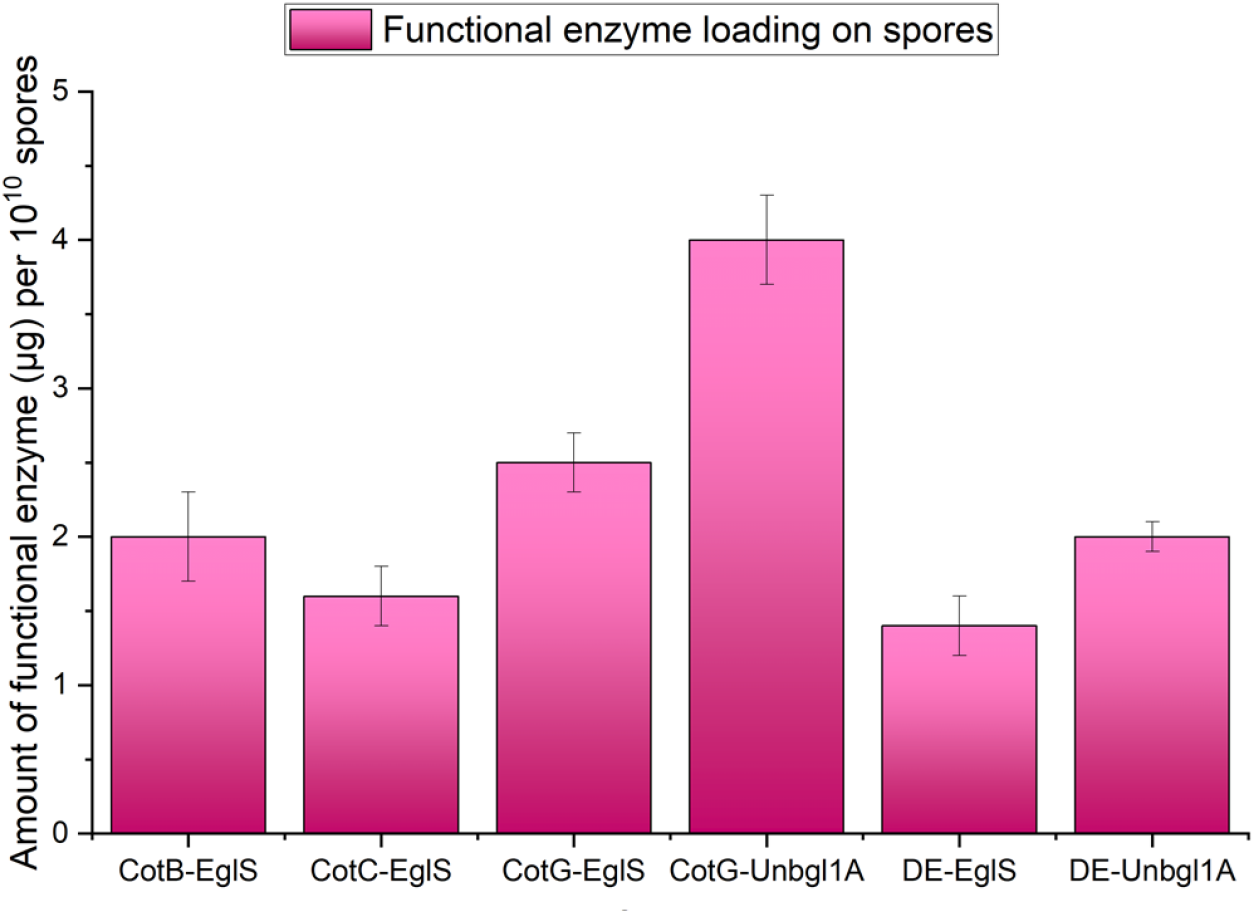
Functional enzyme loading (*µ*g per 10^10^ spores) of single-enzyme EglS and Unbgl1A constructs and the dual-enzyme strain.

### 3.2 Optima and stability of spore-displayed enzymes

The temperature and pH optima were essentially unchanged on display for EglS, while CotG-EglS showed enhanced activity at low pH and elevated temperature, suggesting a stabilising effect of integration into the coat (Figs. 10 and 11). CotG-Unbgl1A showed a shift in temperature optimum to 55°C, compared with 50°C for the free enzyme, with no shift in pH optimum (Figs. 12 and 13).

**Figure 10.**
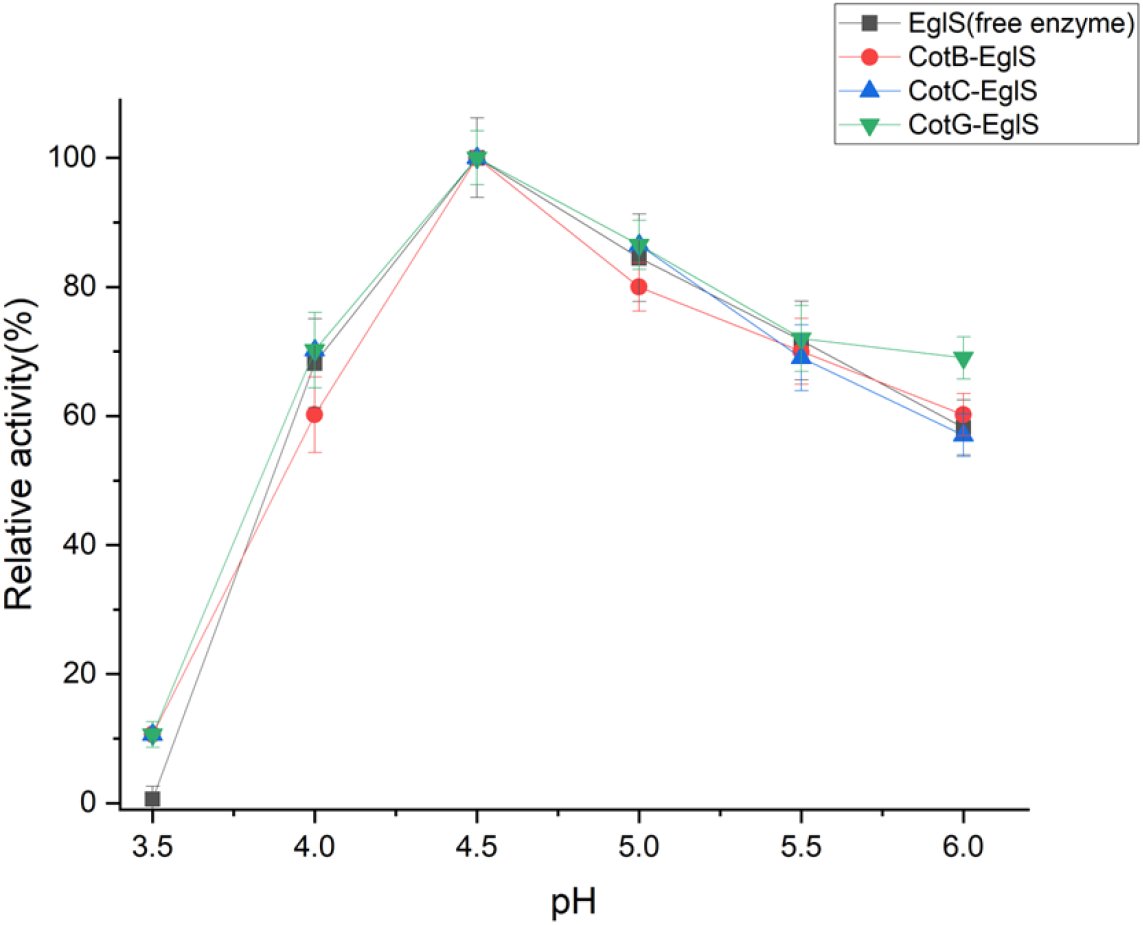
pH optima of free and spore-displayed EglS.

**Figure 11.**
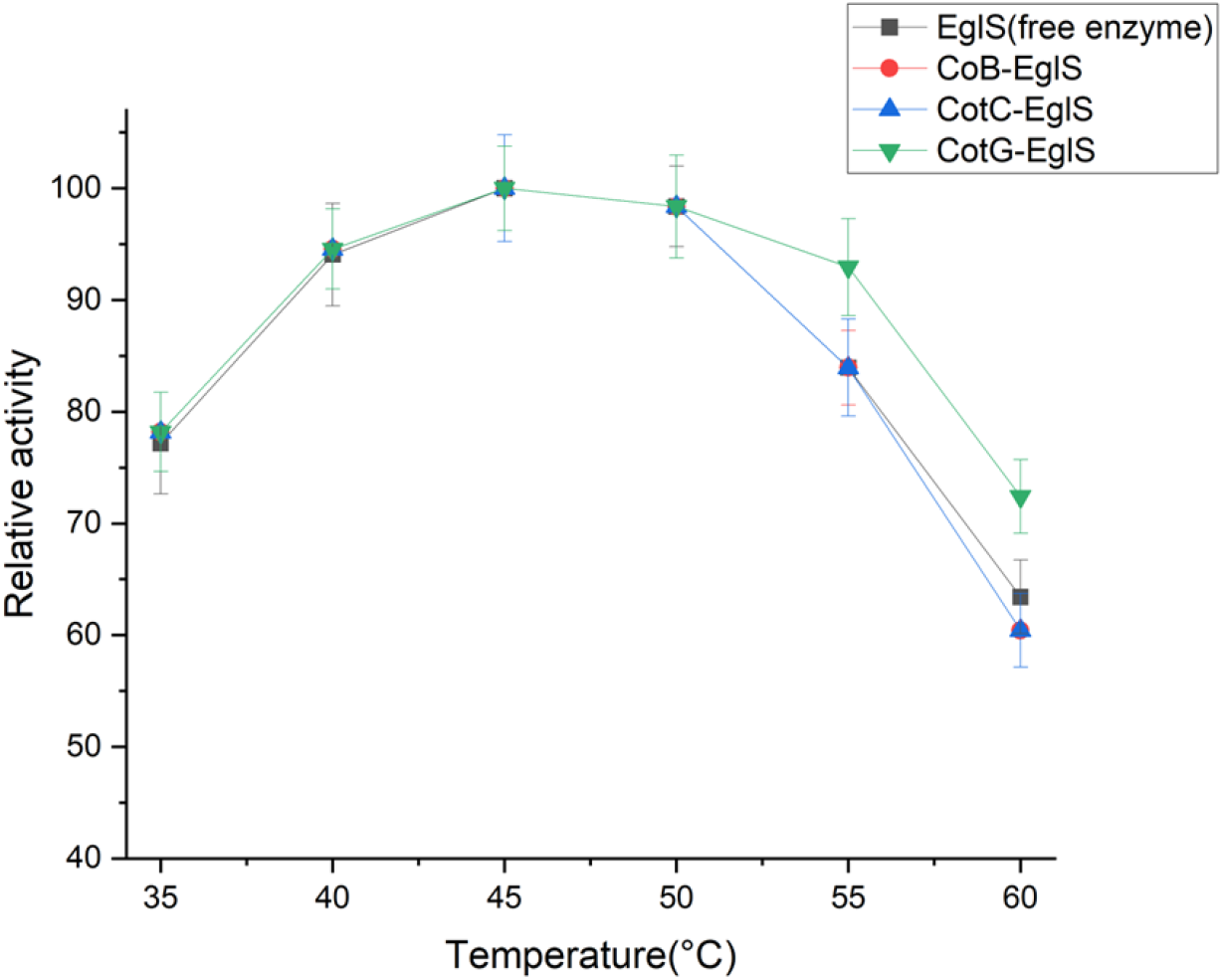
Temperature optima of free and spore-displayed EglS.

**Figure 12.**
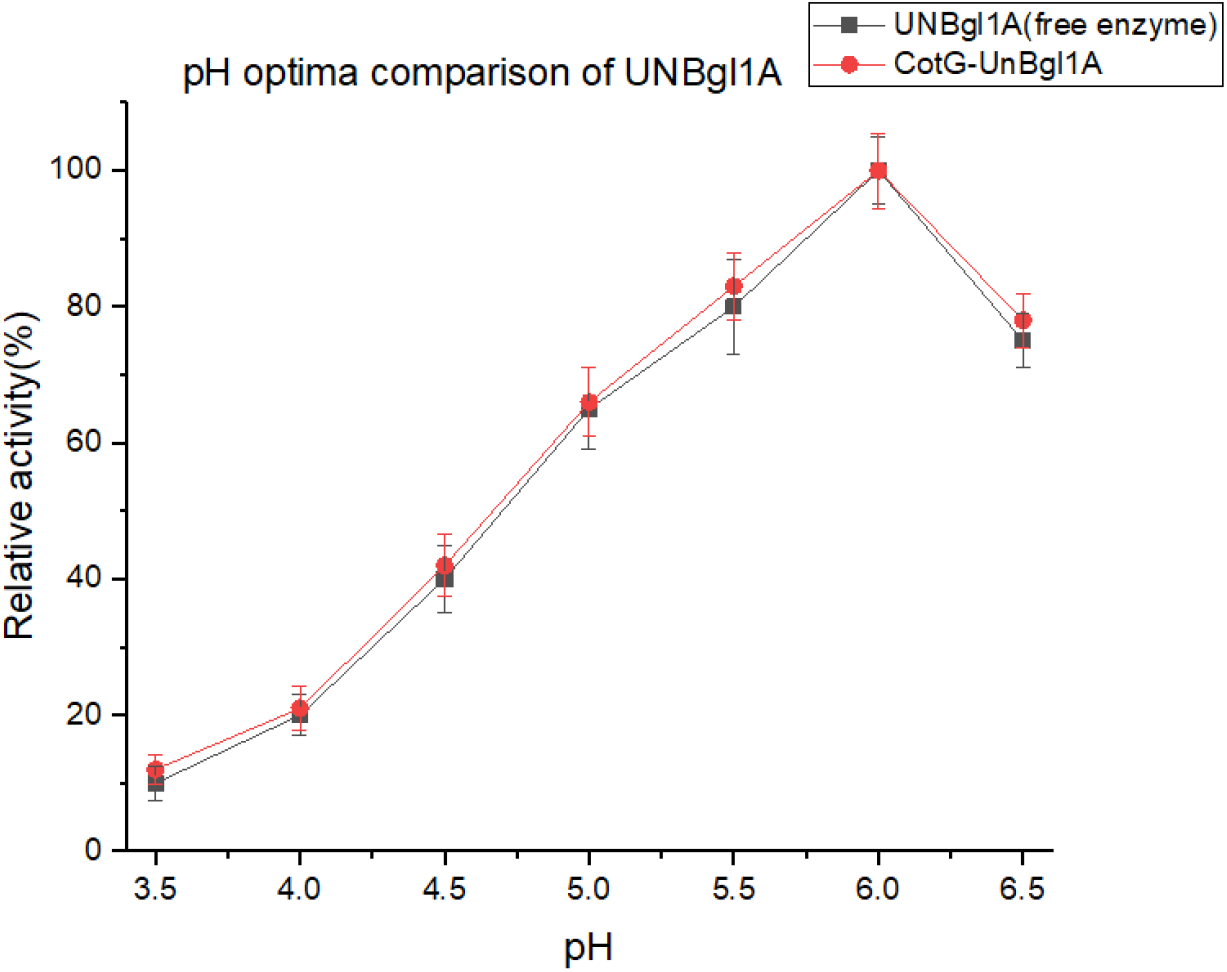
pH optima of free and spore-displayed Unbgl1A.

**Figure 13.**
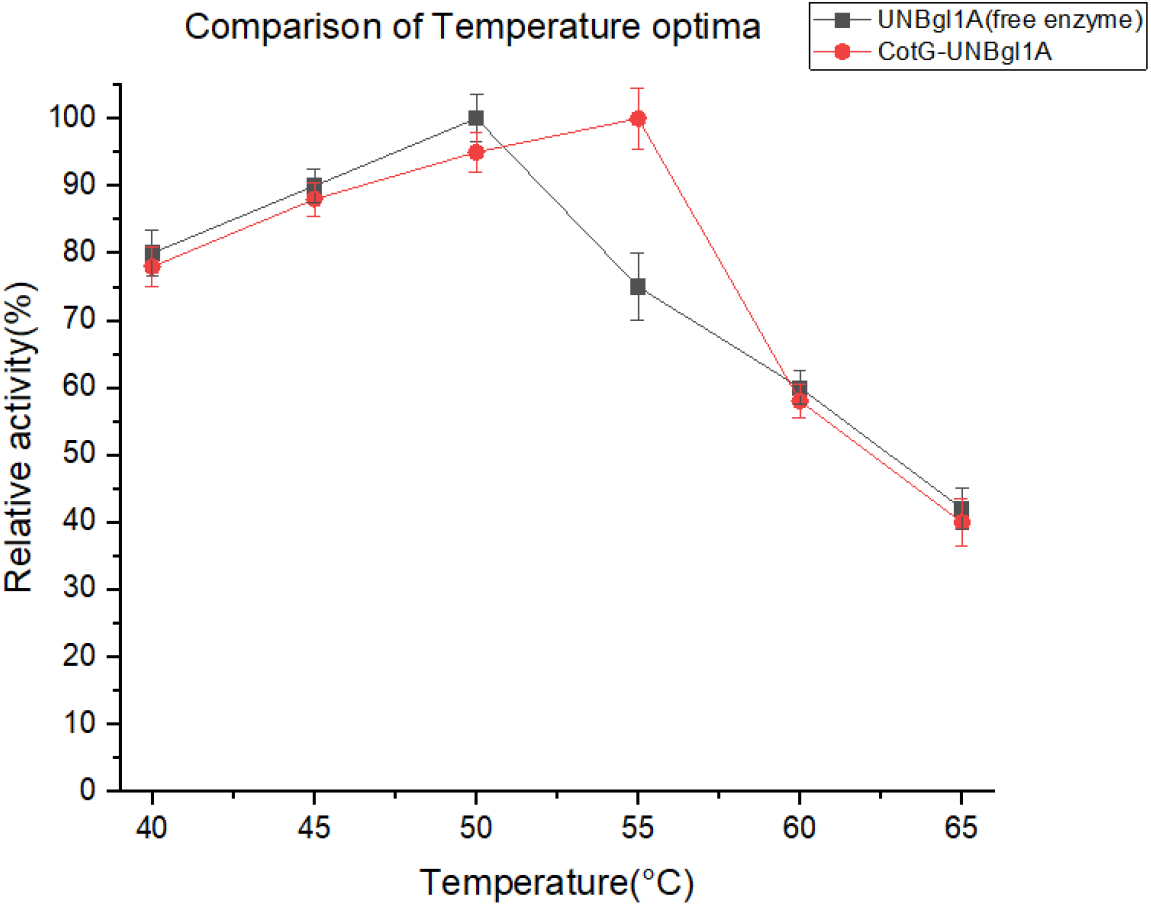
Temperature optima of free and spore-displayed Unbgl1A.

Thermal stability depended on the anchor, with CotG-EglS showing the greatest retention of activity. CotG-EglS retained ~ 46% residual activity after 12 h, against ~ 29% for CotB-EglS and ~ 27% for the free enzyme; CotC-EglS declined rapidly, with no detectable activity after 10 h (Fig. 14). The free enzyme Unbgl1A lost all detectable activity within 6 h, whereas CotG-Unbgl1A retained ~ 30% activity after 8 h (Fig. 15). All three EglS constructs were recyclable: CotG-EglS retained ~ 70% and CotB-EglS ~ 55% activity after four rounds of reuse, whereas CotC-EglS lost activity completely by the third round of reuse (Fig. 16). CotG-Unbgl1A retained ~ 55% activity after four rounds of reuse (Fig. 17). On the basis of this superior thermal stability and recyclability, CotG-EglS was selected as the endoglucanase for the cascade; CotG-Unbgl1A was the only functional *β*-glucosidase construct.

**Figure 14.**
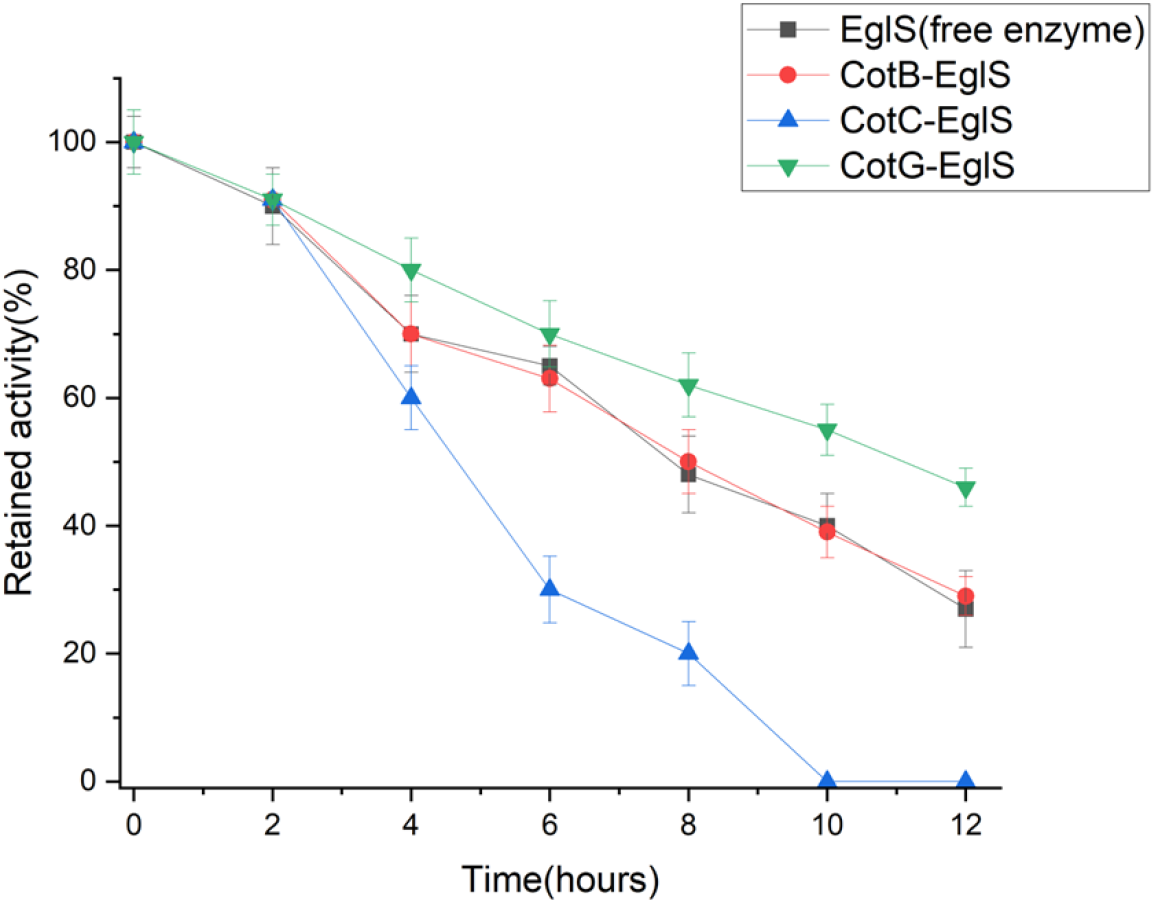
Thermal stability of free and spore-displayed EglS during incubation at 45°C. Residual activity is expressed relative to initial activity (100%).

**Figure 15.**
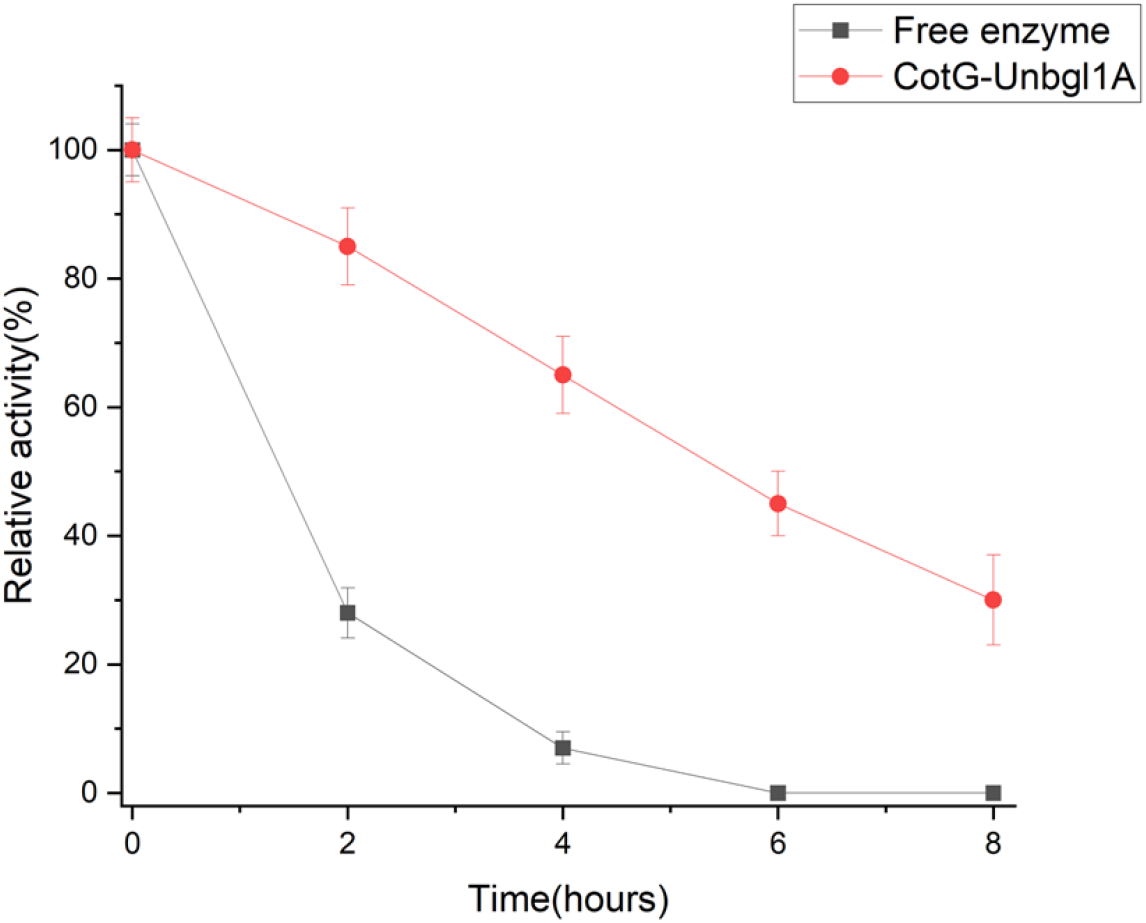
Thermal stability of free Unbgl1A and CotG-Unbgl1A during incubation at 50°C. Residual activity is expressed relative to initial activity (100%).

**Figure 16.**
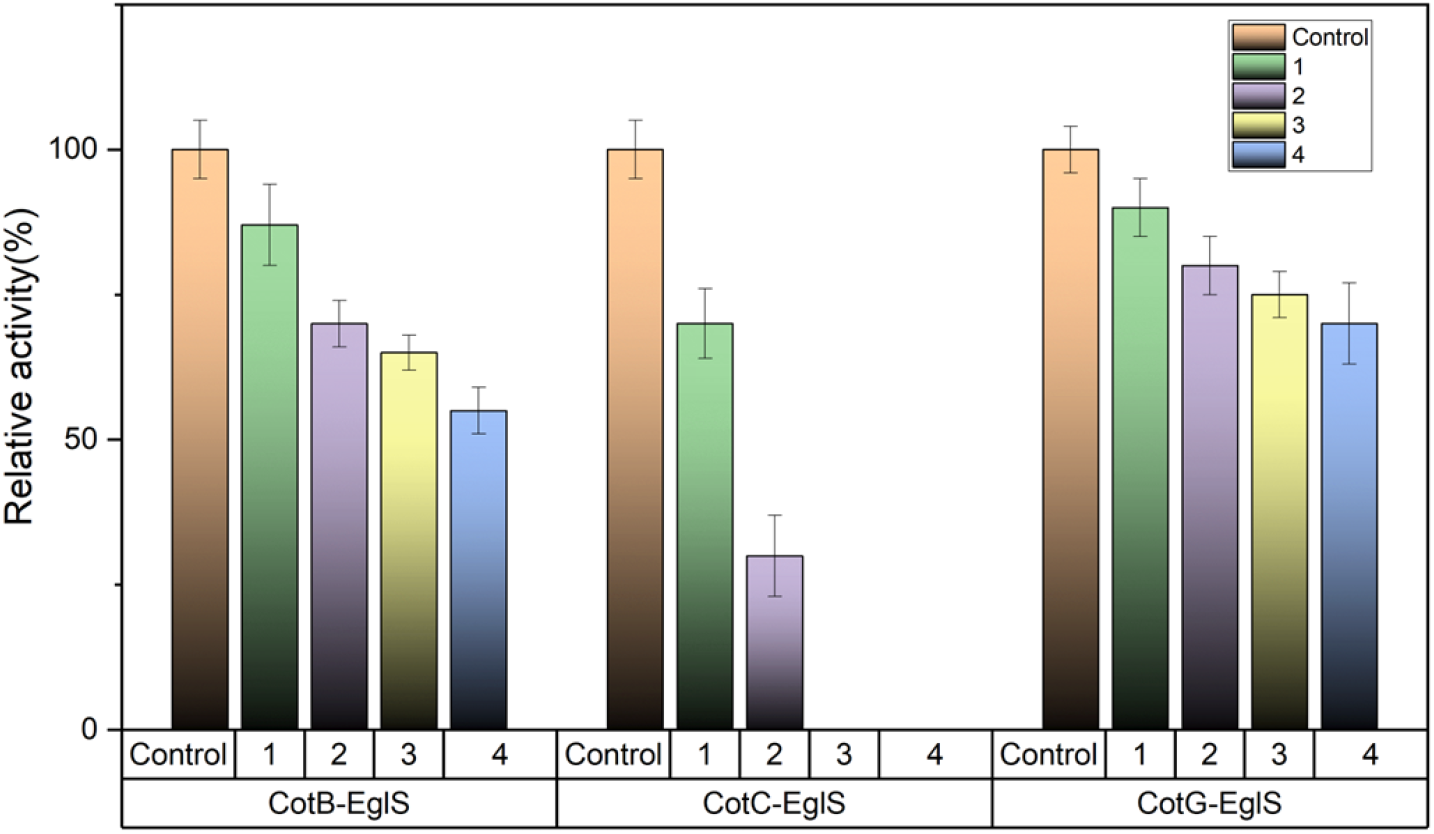
Recyclability of spore-displayed EglS constructs. Control is the first use (100% activity); rounds 1–4 indicate subsequent reuses.

**Figure 17.**
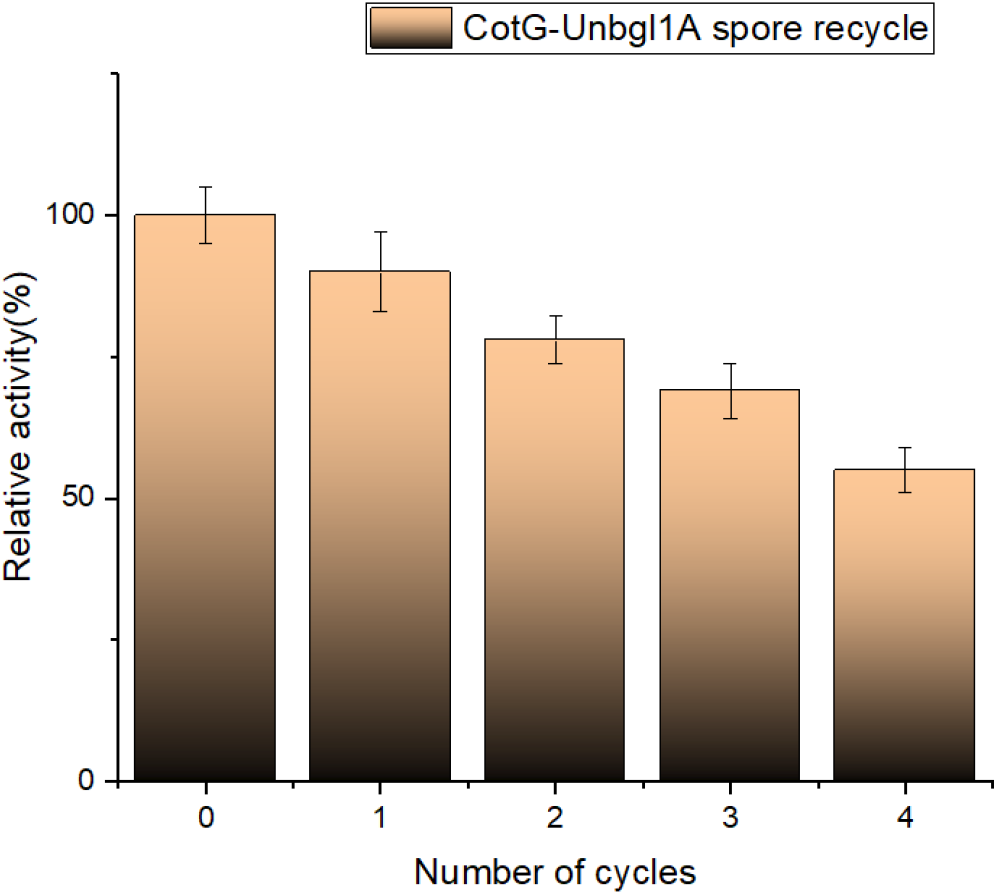
Recyclability of CotG-Unbgl1A. The point labelled 0 is the first use (100% activity); rounds 1–4 indicate subsequent reuses.

### 3.3 Operating conditions for the cascade

Because the two enzymes have different optima, joint operating conditions were determined for the sequential hydrolysis of CMC. Activity increased with pH to a maximum at pH 5 and then dropped sharply, and maximum glucose yield was obtained at 50°C (Figs. 18 and 19). To identify the limiting activity, different EglS:Unbgl1A ratios were tested; doubling EglS (2:1) raised the glucose yield, whereas doubling Unbgl1A had no appreciable effect, identifying the endoglucanase as the rate-limiting activity (Fig. 20). Operating conditions were therefore set to pH 5, 50°C and a 2:1 ratio.

**Figure 18.**
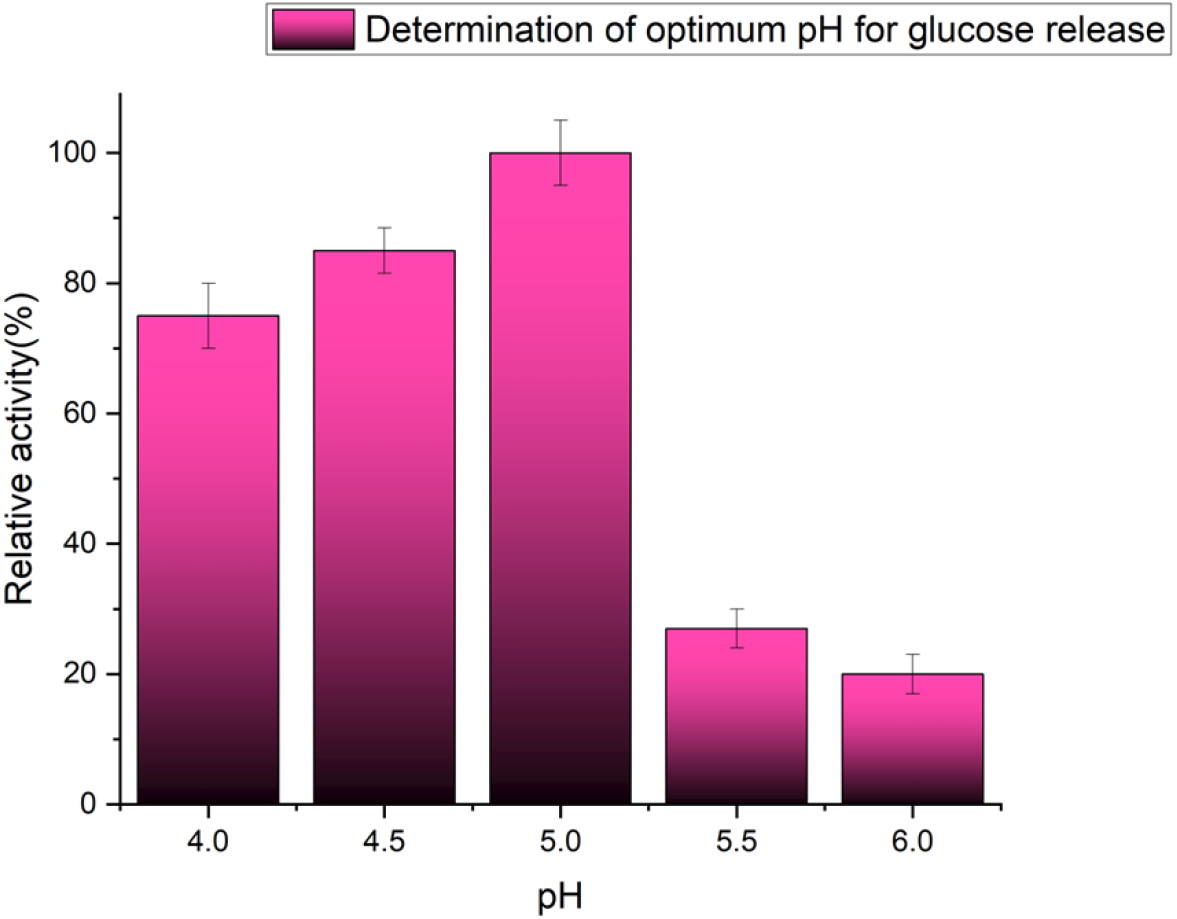
Effect of pH on glucose yield from the two-enzyme cascade.

**Figure 19.**
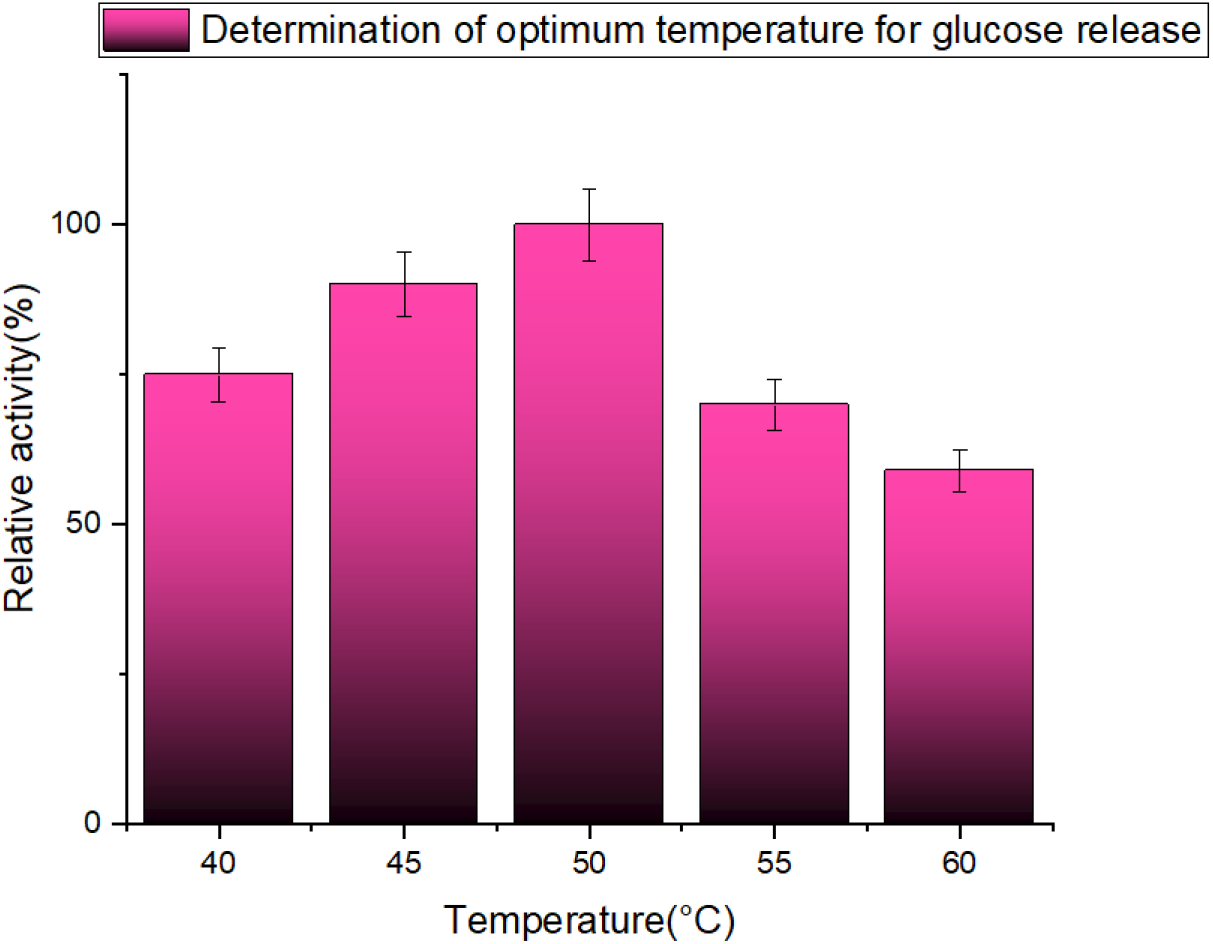
Effect of temperature on glucose yield from the two-enzyme cascade.

**Figure 20.**
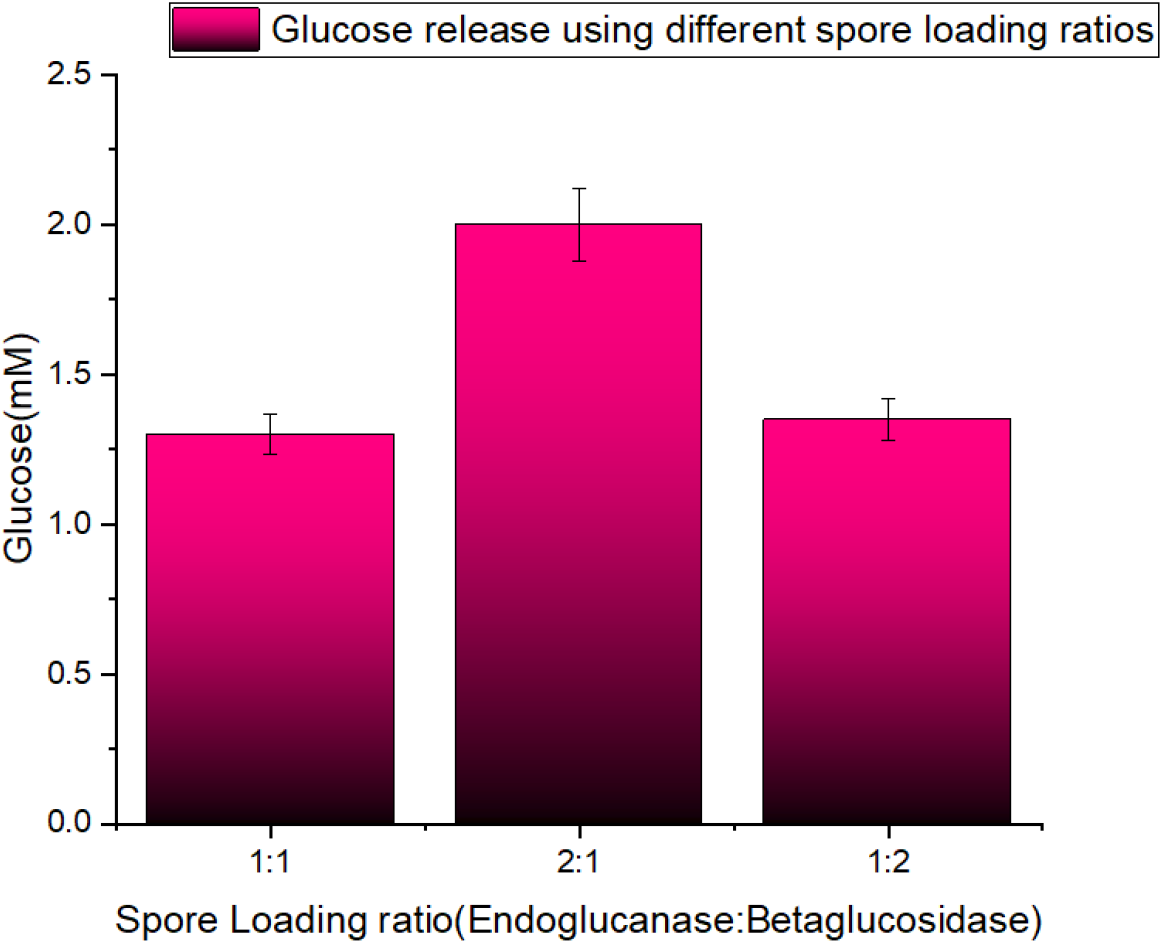
Effect of the EglS:Unbgl1A activity ratio (1:1, 2:1 and 1:2) on glucose yield.

### 3.4 CMC hydrolysis: mixing single-enzyme spores versus dual-enzyme co-display

In the initial comparison under these conditions, the mixture of the two single-enzyme spore strains (CotG-EglS + CotG-Unbgl1A) shown in Fig. 21 produced ~ 2.10 mM glucose in 8 h, exceeding the free-enzyme mixture (~ 1.17 mM) by ~ 79%; the free-enzyme mix showed higher initial activity but tapered off after the fourth hour (Fig. 21). The two single-enzyme constructs and the wild-type control produced little or no glucose individually.

**Figure 21.**
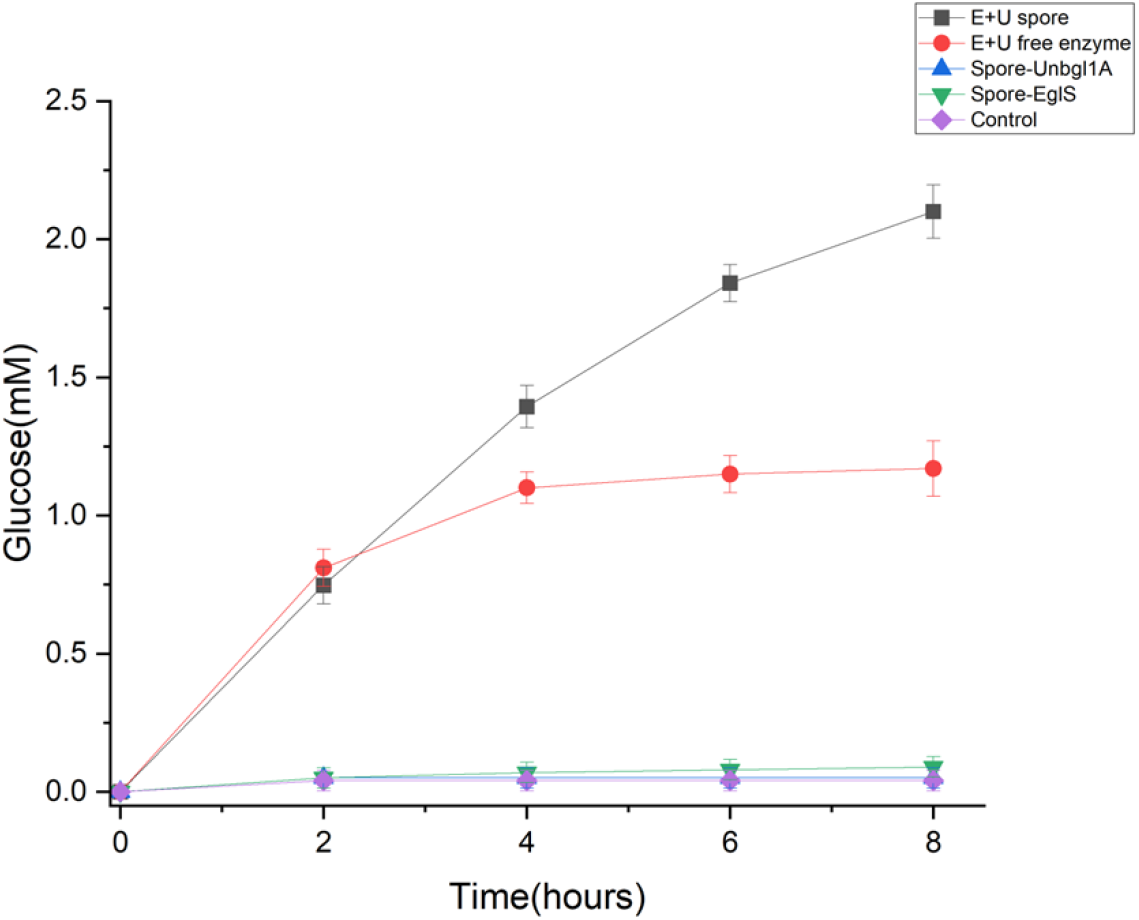
Initial comparison of glucose release from 1% CMC over 8 h at pH 5 and 50°C in 3 mL. Mixed single-enzyme spores (CotG-EglS + CotG-Unbgl1A) and the free-enzyme mixture contained 0.5 U EglS and 0.25 U Unbgl1A. Individual single-enzyme spores and wild-type spores served as controls.

The two constructs were then co-displayed on the single dual-enzyme strain, whose carriage of both integration cassettes was confirmed by selection on both chloramphenicol (pBS1C, integrated at *amyE*) and spectinomycin (pBS4S, integrated at *thrC*). Because the strain is a single clonal isolate carrying both single-copy chromosomal cassettes, every spore inherits both cassettes, and spores from this strain are therefore expected to express both fusions during sporulation; co-display of EglS and Unbgl1A on the same spore thus follows from the construction of the strain rather than being inferred from the reaction. After construction of the dual-enzyme strain, the comparison was repeated in a separate experiment with all three preparations, each containing 0.5 U EglS. The dual-enzyme strain produced 1.57 mM glucose, outperforming the free-enzyme mix (1.35 mM) by ~ 16% and confirming that both displayed activities are catalytically functional together in the cascade (Fig. 22), with both enzymes also independently quantified on the dual-strain spores (Fig. 9). However, the yield from the dual-enzyme strain was ~27% lower than that from the mixed single-enzyme spores.

**Figure 22.**
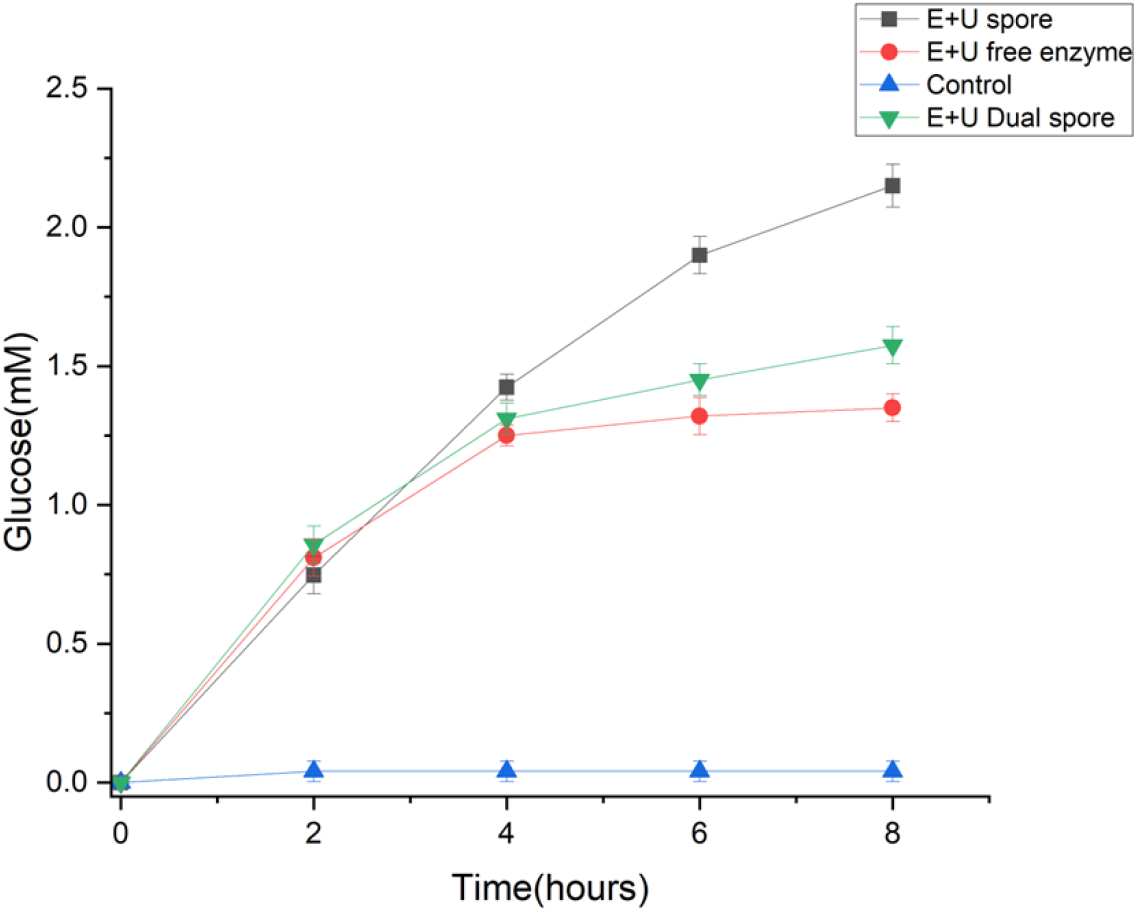
Separate comparison performed after construction of the dual-enzyme strain: glucose release from 1% CMC over 8 h at pH 5 and 50°C in 3 mL. EglS activity was matched at 0.5 U in the dual-enzyme strain, mixed single-enzyme spores and free-enzyme mixture. Corresponding Unbgl1A activities were 0.3, 0.25 and 0.25 U, respectively.

## 4 Discussion

To our knowledge, this study reports the first case of two distinct, catalytically active enzymes co-displayed on a single *Bacillus subtilis* spore surface. A recent study demonstrated a two-enzyme cascade on spores, but the enzymes were displayed on separate spore populations and then mixed, so the two activities resided on different spores rather than being co-displayed on the same spore (Spannenkrebs et al., 2026); in the dual strain reported here, by contrast, both fusion cassettes are chromosomally integrated in one clonal isolate, so every spore carries both and is expected to display them. Earlier reports of more than one passenger per spore differ in important ways: a single enzyme, trehalose synthase, was displayed using two cot partners (CotC-TreS and CotG-TreS) integrated into the same strain to increase the number of displayed molecules (Liu et al., 2019), and GFP and RFP were co-displayed on a single spore, but that study focused on overexpression at varying temperatures, with no data on stability or activity (Petrillo et al., 2020). Here, CotG-EglS and CotG-Unbgl1A were selected for co-display because CotG gave the most stable and recyclable endoglucanase construct and was the only functional anchor for Unbgl1A.

Immobilisation can restrict enzyme conformational flexibility, which may contribute to improved stability (Secundo, 2013). This offers a possible explanation for the enhanced thermal stability observed here, although conformational changes were not measured. Changes in catalytic optima and improvements in thermal and pH stability have also been reported for other *Bacillus subtilis* spore-displayed enzymes (Shi et al., 2025; Li et al., 2024; Mingmongkolchai and Panbangred, 2019; Sheng et al., 2017). The approximately 5°C increase in the temperature optimum of CotG-Unbgl1A may likewise reflect changes in enzyme conformation or its local environment following surface display.

The lower glucose yield of the dual-enzyme strain cannot be attributed to a lower starting EglS activity, since this was matched at 0.5 U in all three preparations. The mixed single-enzyme spores sustained glucose production for longer, suggesting that differences in operational stability contributed to the yield difference. Germination may have contributed to activity loss in the dual-enzyme preparation, as vegetative cells were observed by microscopy in the reaction mixture. Spore coat formation depends on the ordered assembly and interactions of coat proteins (McKenney et al., 2013; Henriques and Moran Jr., 2007; Krajčíková et al., 2017). Overexpression of the two CotG fusion proteins during sporulation may interfere with spore coat assembly in the dual-enzyme strain. Defects in the assembly of spore protective layers have been linked to premature, GerA-mediated germination during sporulation in *Bacillus subtilis* (Ramírez-Guadiana et al., 2017), although whether the fusion proteins caused such defects in this study was not determined. That study concerns germination during spore development, so it does not directly explain germination during our enzyme assay. Prolonged incubation at elevated temperature (Luu et al., 2015) and the presence of glucose as a potential co-germinant (Fan et al., 2024) are additional factors that may have contributed. The relative contributions of enzyme inactivation, germination and fusion behaviour to activity loss were not determined. Previous spore-display studies have reduced germination-associated losses using strains lacking the cortex-lytic genes *sleB* and *cwlJ* (Liu et al., 2019; Sun et al., 2020; Chirakkal et al., 2002; Heffron et al., 2009).

Industrially, multi-enzyme synthesis is often carried out by co-culturing strains that house distinct enzymes (Zhang and Wang, 2016) or by metabolic division of labour in engineered consortia (Duncker et al., 2021). Mixing single-enzyme spore strains allows the enzyme ratio to be adjusted between batches, whereas the dual-enzyme strain has a fixed, strain-determined ratio. This flexibility has also been noted for other spore-based cascades (Spannenkrebs et al., 2026); it is a practical advantage distinct from the difference in sustained activity observed here. A systematic comparison of the same enzyme across the three commonly used outer-coat anchors, with matched activity, stability, recyclability and loading data, has also been lacking; the closest prior example displayed a keratinase on CotB and CotG and likewise found CotG to be the more effective partner (Wang et al., 2024). EglS was active with all three anchors and could be quantified using a simple colourimetric assay. These properties make it a promising reporter for comparing anchor performance, although reporter development was not the primary aim of this study.

## 5 Conclusion

Two distinct cellulolytic enzymes, an endoglucanase and a *β*-glucosidase, were functionally co-displayed on a single *Bacillus subtilis* spore, and the resulting strain catalysed the two-step hydrolysis of CMC to glucose. CotG was the most effective anchor for EglS and the only functional anchor for Unbgl1A, with improved thermal stability relative to the free enzymes and activity retained over repeated reuse. However, mixing separate single-enzyme spore strains gave a higher glucose yield than the single dual-enzyme strain and allowed the enzyme ratio to be tuned to the rate-limiting endoglucanase. For CMC hydrolysis under the conditions tested, mixing separately displayed single-enzyme spores gave higher glucose yields and offered greater flexibility in setting enzyme proportions than co-display on one spore, while co-display remains attractive where a fixed, strain-determined enzyme ratio is acceptable.

## Acknowledgements

The authors thank the current and former members of the Protein Engineering Lab for helpful discussions and technical assistance, and the Department of Chemical Engineering, Indian Institute of Technology Bombay, for providing the research facilities used in this work.

## Statements and Declarations

### Author contributions

R.P.J. and S.B.N. conceived and designed the study. R.P.J. performed the experiments, analysed the data, and wrote the manuscript. S.B.N. supervised the project, acquired funding, and reviewed and edited the manuscript. Both authors read and approved the final manuscript.

### Ethics approval

Not applicable. This study did not involve human participants or animals.

### Consent to participate / Consent for publication

Not applicable.

### Data availability

The data supporting the findings of this study are available from the corresponding author upon reasonable request.

## References

Borges DG, Junior AB, Farinas CS, et al (2014) Enhanced saccharification of sugarcane bagasse using soluble cellulase supplemented with immobilized β-glucosidase. Bioresource Technology 167:206–213. 10.1016/j.biortech.2014.06.021

Chirakkal H, O’Rourke M, Atrih A, et al (2002) Analysis of spore cortex lytic enzymes and related proteins in Bacillus subtilis endospore germination. Microbiology 148(8):2383–2392. 10.1099/00221287-148-8-2383

Duncker KE, Holmes ZA, You L (2021) Engineered microbial consortia: strategies and applications. Microbial Cell Factories 20(1):211. 10.1186/s12934-021-01699-1

Fan L, Zhang Y, Ismail BB, et al (2024) Bacillus spore germination: mechanisms, identification, and antibacterial strategies. Critical Reviews in Food Science and Nutrition 64(30):11146–11160

Harwood CR, Cutting SM (eds) (1990) Molecular Biological Methods for Bacillus. John Wiley, Chichester

Heffron JD, Orsburn B, Popham DL (2009) Roles of germination-specific lytic enzymes CwlJ and SleB in Bacillus anthracis. Journal of Bacteriology 191(7):2237–2247. 10.1128/JB.01506-08

Henriques AO, Moran Jr. CP (2007) Structure, assembly, and function of the spore surface layers. Annual Review of Microbiology 61(1):555–588. 10.1146/annurev.micro.61.080706.093224

Iwanicki A, Piątek I, Stasiłojć M, et al (2014) A system of vectors for Bacillus subtilis spore surface display. Microbial Cell Factories 13:30. 10.1186/1475-2859-13-30

Krajčíková D, Forgáč V, Szabo A, et al (2017) Exploring the interaction network of the Bacillus subtilis outer coat and crust proteins. Microbiological Research 204:72–80. 10.1016/j.micres.2017.07.002

Lehmann M, Herrmann M (2025) Enzyme-displaying spores as a novel strategy for mixed fiber textile recycling. Frontiers in Systems Biology 5:1603731. 10.3389/fsysb.2025.1603731

Li S, He L, Shi N, et al (2024) Display of lignin peroxidase on the surface of Bacillus subtilis. Applied Biochemistry and Biotechnology 196(10):6849–6863

Liu H, Yang S, Wang X, et al (2019) Production of trehalose with trehalose synthase expressed and displayed on the surface of Bacillus subtilis spores. Microbial Cell Factories 18:1–13. 10.1186/s12934-019-1068-0

Luu S, Cruz-Mora J, Setlow B, et al (2015) The effects of heat activation on Bacillus spore germination, with nutrients or under high pressure, with or without various germination proteins. Applied and Environmental Microbiology 81(8):2927–2938. 10.1128/AEM.04257-14

McKenney PT, Driks A, Eichenberger P (2013) The Bacillus subtilis endospore: assembly and functions of the multilayered coat. Nature Reviews Microbiology 11(1):33–44. 10.1038/nrmicro2921

Mingmongkolchai S, Panbangred W (2019) Display of Escherichia coli phytase on the surface of Bacillus subtilis spore using CotG as an anchor protein. Applied Biochemistry and Biotechnology 187:838–855. 10.1007/s12010-018-2855-7

Nath P, Dhillon A, Kumar K, et al (2019) Development of bi-functional chimeric enzyme (CtGH1-L1-CtGH5-F194A) from endoglucanase (CtGH5) mutant F194A and β-1, 4-glucosidase (CtGH1) from Clostridium thermocellum with enhanced activity and structural integrity. Bioresource Technology 282:494–501

Nguyen NNY, Duong L, Nguyen TT, et al (2025) Oral administration of Bacillus subtilis spores expressing Staphylococcus aureus IsdB induces mucosal immune responses in mice. Biotechnology Letters 47(4):75. 10.1007/s10529-025-03620-5

Petrillo C, Castaldi S, Lanzilli M, et al (2020) The temperature of growth and sporulation modulates the efficiency of spore-display in Bacillus subtilis. Microbial Cell Factories 19:1–10. 10.1186/s12934-020-01446-6

Radeck J, Kraft K, Bartels J, et al (2013) The Bacillus BioBrick Box: generation and evaluation of essential genetic building blocks for standardized work with Bacillus subtilis. Journal of Biological Engineering 7:1–17. 10.1186/1754-1611-7-29

Rahman MS, Fernando S, Ross B, et al (2018) Endoglucanase (EG) activity assays, vol 1796, Humana Press Inc., pp 169–183. 10.1007/978-1-4939-7877-9_13

Ramírez-Guadiana FH, Meeske AJ, Wang X, et al (2017) The Bacillus subtilis germinant receptor GerA triggers premature germination in response to morphological defects during sporulation. Molecular Microbiology 105(5):689–704. 10.1111/mmi.13728

Secundo F (2013) Conformational changes of enzymes upon immobilisation. Chemical Society Reviews 42(15):6250–6261. 10.1039/c3cs35495d

Sheng S, Jia H, Topiol S, et al (2017) Engineering CotA laccase for acidic pH stability using Bacillus subtilis spore display. Journal of Microbiology and Biotechnology 27(3):507–513. 10.4014/jmb.1608.08026

Shi N, Cheng D, Chen Y, et al (2025) Improvement in lignin peroxidase oxidative stability via surface display by Bacillus subtilis. Biotechnology Letters 47(4):86. 10.1007/s10529-025-03625-0

Spannenkrebs JB, Klau LJ, Karava M, et al (2026) Spore immobilized enzymes for the multi-step synthesis of cellobiose. Microbial Cell Factories 25(1):43. 10.1186/s12934-026-02943-w

Sun R, Zhang M, Chen H, et al (2020) Germination-arrest Bacillus subtilis spores as an oral delivery vehicle of grass carp reovirus (GCRV) Vp7 antigen augment protective immunity in grass carp (Ctenopharyngodon idella). Genes 11(11):1351. 10.3390/genes11111351

Wang H, Wang Y, Yang R (2017) Recent progress in Bacillus subtilis spore-surface display: concept, progress, and future. Applied Microbiology and Biotechnology 101:933–949. 10.1007/s00253-016-8080-9

Wang Z, Yan M, Saeed M, et al (2024) The flexible linker and CotG were more effective for the spore surface display of keratinase KERQ7. World Journal of Microbiology and Biotechnology 40(1):35

Zhang H, Wang X (2016) Modular co-culture engineering, a new approach for metabolic engineering. Metabolic Engineering 37:114–121. 10.1016/j.ymben.2016.05.007

